# Do we parse the background into separate streams in the cocktail party?

**DOI:** 10.1101/2022.02.21.480990

**Authors:** Orsolya Szalárdy, Brigitta Tóth, Dávid Farkas, Gábor Orosz, István Winkler

**Author notes:** Corresponding author: Orsolya Szalárdy, Address: H-1089 Budapest, Nagyvárad tér 4.

## Abstract

In the cocktail party situation, people with normal hearing usually follow a single speaker among multiple concurrent ones. However, there is no agreement in the literature as to whether the background is segregated into multiple streams/speakers. The current study varied the number of concurrent speech streams and investigated target detection and memory for the contents of a target stream as well as the processing of distractors. A male-spoken target stream was either presented alone (single-speech), together with one male-spoken (one-distractor), or a male- and a female-spoken distractor (two-distractor). Behavioral measures of target detection and content tracking performance as well as target- and distractor detection related ERPs were assessed. We found that the detection sensitivity and the target N2b amplitude decreased whereas the P3b amplitude increased from the single-speech to the concurrent speech streams conditions. Importantly, the behavioral distractor effect differed between the conditions with one- vs. two-distractor (distraction by the female speaker was lower than that of the male speaker in either condition) and the target N2b elicited in the presence of two distractors was significantly smaller than that elicited in the presence of one distractor. Further, the voltage in the N2b time window significantly differed between the one- and two-distractor conditions for the same (M2) speaker. These results show that speech processing was different in the presence of one vs. two distractors, and thus, the current data suggest that the two background speech streams were segregated from each other.

## Introduction

In everyday environments, we often attend to speech in the presence of multiple other speech streams (termed the “cocktail party” situation; Cherry, 1953). Typically, the listener’s goal is to follow the content of one speech stream while a speech from other talkers may distract him/her. People with normal hearing usually manage this situation (see, e.g. Bregman, 1990; Wood & Cowan, 1995). To this end, the auditory system must decompose the mixture of sounds into meaningful streams (auditory scene analysis; Bregman, 1990) and select the one with the behaviorally relevant information (selective attention; Astheimer & Sanders, 2009; Best et al., 2008). At the same time, processing the irrelevant streams should be suppressed to some degree in order to conserve capacities and prevent distraction (Ihlefeld & Shinn-Cunningham, 2008; Szalárdy et al., 2019; Szalárdy, Tóth, Farkas, Orosz, et al., 2020b). Some studies showed that the auditory system can use a foreground-background solution with sounds in the acoustic background not being separated to further streams (e.g., Brochard et al., 1999; for a review, see Snyder & Alain, 2007). However this might not be always the case, for instance when some of the background streams contain distinct auditory features (as suggested by Cusack et al., 2004; Winkler et al., 2012). In the current study, we tested whether two non-target speech streams from speakers of different gender are segregated in the presence of a third (target) speech stream. To this end, we assessed performance in target detection and extracting and storing content information from the target speech stream and the processing of distractors presented in one vs. two distractor speech streams using behavioral measures and event-related brain potentials (ERPs).

Speech processing in the presence of other concurrent sound streams has been the target of several studies (for a recent review, see Bronkhorst, 2015). These studies mostly reported higher processing demand in the presence of concurrent speech compared to that with a single speech stream, as indicated by both behavioral and neural measures (see, e.g. Lambrecht et al., 2011). When speech was used for the distractor, stronger masking and reduced target detection performance were observed for the target speech stream compared to spectrally matched noise distractors (Kidd et al., 2005). Speech streams to be suppressed may lead to allocation errors through information masking. For example, when listeners were instructed to detect words in the target speech stream, there was a significant chance of reporting words from the distractor (masker) speech stream (Kidd et al., 2005; Wightman & Kistler, 2005). In our previous study (Szalárdy et al., 2019), listeners heard two concurrent speech streams, and they were instructed to detect numerals in the target stream. We found reduced detection sensitivity (d’), decreased hit, and increased false alarm rates with increased information masking. Furthermore, informational masking has been shown to influence the neural representation of the target speech (Szalárdy et al., 2019; Yuriko Santos Kawata et al., 2020).

The issue of auditory foreground-background decomposition has also been addressed by several experimental and theoretical papers (Siegenthaler & Barr, 1967; Teki et al., 2011; Tóth et al., 2016). Many of these papers suggest that when the auditory scene is segregated into streams, one of the streams can be consciously perceived, forming the auditory foreground while the rest of the auditory scene falls outside conscious perception, forming the background. This notion is, for example, supported by studies measuring the mismatch negativity (MMN, an event-related potential elicited by violations of acoustic regularities; for a recent review, see 2020), as some studies found MMN only to deviants violating regularities of the currently consciously experienced sound sequence (i.e., the foreground; Rahne et al., 2007; Sussman et al., 1998; Winkler et al., 2006).

Somewhat less is known about the extent to which the background stream is processed. Some studies showed that occasionally, sounds from the background may intrude into consciousness, for instance, some unexpected or personally relevant acoustic event (see, e.g. Micheyl et al., 2007), but not regularities, *per se* (Southwell et al., 2017). Furthermore, there is also evidence showing that violations of some regularities are detected also within the background stream (Szalárdy, Winkler, et al., 2013) and, in general, stream segregation can occur outside the focus of attention (Bregman, 1990; Sussman et al., 2007). Thus, the question remains, whether sounds outside the focus of attention form an unsegregated background or the processing received by sounds outside the focus of attention includes stream segregation. Winkler and colleagues (2012) described the alternatives, arguing for a full segregation model (Mill et al., 2013). Here we test this possibility for concurrent speech streams.

Event-related potentials (ERP) were measured, because they allow one to study processes of target detection, attentional selection, working memory, and distraction. Target auditory events (including speech stimuli) typically elicit two successive ERP components, the N2b and the P3b (Näätänen et al., 1982; Polich, 2007; Polich & Herbst, 2000; Ritter et al., 1983). The N2b is a negative potential reaching maximal amplitude at around 200 ms from stimulus onset with a typical centro-parietal scalp distribution. In contrast to other subcomponents from the N2 family, N2b typically appears after a detected target event and has been associated with stimulus classification (Näätänen, 1990; Ritter et al., 1979). Studies have found that the amplitude of N2b is modulated by selective attention (Michie, 1984) and stream segregation (Szalárdy, Bőhm, et al., 2013). The N2b is often followed by the P3b, which is a positive potential usually peaking between 300 and 400 ms from stimulus onset and with a parietally dominant scalp distribution (Conroy & Polich, 2007; Polich, 2003). P3b has been associated with context updating (Donchin & Coles, 1988), categorization, and later evaluation of the target stimulus (Nasman & Rosenfeld, 1990). The P3b has been shown to reflect the interaction between selective attentional processes and working memory (Polich & Herbst, 2000). This component has been selectively modulated by informational (and energetic) masking in our previous experiment (Szalárdy et al., 2019), resulting in reduced P3b when poorer allocation of attention could be assumed. Both components appear with larger amplitude with increased cognitive demand (Isreal et al., 1980; Polich, 2007). For non-target surprising events, another subcomponent from the P3 group is elicited, the P3a or novelty P3 (Polich, 2007). In a previous study, this component was elicited by target-like events appearing within a non-target speech stream delivered concurrently to the target speech stream (Szalárdy, Tóth, Farkas, Orosz, et al., 2020b). In the current study, N2b and P3b will be used to assess the effects of the manipulations on target detection and processing of distractors.

A continuous target speech stream was presented to the participants alone (single-speech condition) or in the presence of one or two continuous distractor speech stream(s) (one-distractor and two-distractor condition, respectively). Participants were instructed to detect numerals in the target stream by pressing a reaction key (target detection task). They were also asked to follow the target speech and to answer questions based on the information presented in it (recognition task). The target stream was delivered by a male speaker. The first distractor stream was spoken by a different male speaker (both in the one- and the two-distractor condition) with a female speaker delivering the third speech stream in the two-distractor condition. We hypothesized that performance (both target detection and recognition performance) will be lower in the conditions with distractor speech streams compared to the single-speech condition. Concurrently, based on previous studies showing that the amplitude of the N2b/P3b amplitudes to target events increase with increasing task demand (Isreal et al., 1980; Polich, 2007; Szalárdy, Tóth, Farkas, Hajdu, et al., 2020), we hypothesized that the N2b/P3b elicited by target numerals will be larger in the presence of distractor speech compared to the single-speech condition. By using two distractor speech streams delivered by speakers of different gender, we aimed to provide acoustically sufficiently distinct stimuli to promote segregation of the two background streams. If the two non-target streams are segregated, then one should expect significant performance and target-related ERP-amplitude decrease from the one- to the two-distractor condition due to the additional processes required for segregating the two non-target streams. In contrast, small or no performance and ERP differences between the one- and the two-distractor condition would suggest a predominantly foreground/background solution of the two-distractor condition by the auditory system.

## Methods

### Participants

Native Hungarian speakers (N = 29; 11 males; age: M = 21.97 years, SD = 2.04; 26 right-handed) participated in the study for modest financial compensation. None of the participants had a history of psychiatric or neurological symptoms. All participants had pure-tone thresholds of <25 dB in the 250 Hz - 4 kHz range, with <10 dB difference between the two ears. Data from two participants were excluded from the final analysis due to the loss of the EEG triggers for sound onset. Data from four paritipants were excluded tue to extensive artefacts and bad signal-to noise ratio. Thus, data from 23 participnats were analysed (8 male, 15 female, mean age: 21.91 years, SD: 2.23, 21 right-handed). Written informed consent was obtained from all participants, and modest financial compensation was given for participating in the experiment. The study was approved by the United Hungarian Ethical Committee for Research in Psychology (EPKEB), and it was in full compliance with the World Medical Association Helsinki Declaration and all applicable national laws.

### Stimuli

Speech recordings of approximately 6 minutes duration were used as stimuli (soundtracks recorded at 48 kHz with 32-bit resolution, mean duration: 355.33 s, SD: 12.28, mean word number per stream: 636.41, SD: 84.87; mean number of phonemes per word: 6.48, SD: 0.29). Hungarian, emotionally neutral informative articles of news websites were delivered by professional actors (two male and one female speaker) recorded at 48 kHz with 32-bit resolution in the same room where the experiment was conducted.

All articles were reviewed by a dramaturge checking for correct syntax and natural flow of the text. The recorded speech was edited by a professional radio technician. The average RMS of the sound recordings was equalized to −32dBfs by VST-based attenuation after applying either −20 dB or −15 dB C3 compressors, depending on the dynamics of the actor reading. Audio recordings were presented by Matlab R2014a software (Mathworks Inc.) with Psychtoolbox 3.0.10 on two Intel Core i5 PCs with ESI Julia 24-bit 192 kHz sound cards connected to Mackie MR5 mk3 Powered Studio Monitor loudspeakers. The speech streams were presented with a fixed loudness level of ~70 dB SPL. Speech recordings were delivered from the same position as they were recorded in order to recreate the reverberation effects of the recording situation. Thus, room acoustics effects did not differ between recording and the experimental setup. Each loudspeaker corresponded to one speaker.

In three experimental conditions, one, two, or three speech streams were presented concurrently (see Figure 1A for a schematic illustration). Speech from the left loudspeaker (a male speaker’s speech) was designated as the target of the task (target stream). When delivered, the other stream(s) served as the distractor(s): the other male speaker’s speech in the two-stream condition, whereas both the other male speaker’s and the female speaker’s speech in the three-stream condition. The spatial arrangement of the distractor streams was also constant throughout the experiment: the male actor’s speech was presented from the right, while the female actor’s speech from the central loudspeaker. The starting times of the audio playbacks for concurrent speech streams were synchronized by a microcontroller ensuring that all speech segments started within a 6 ms timeframe.

**Figure 1.**
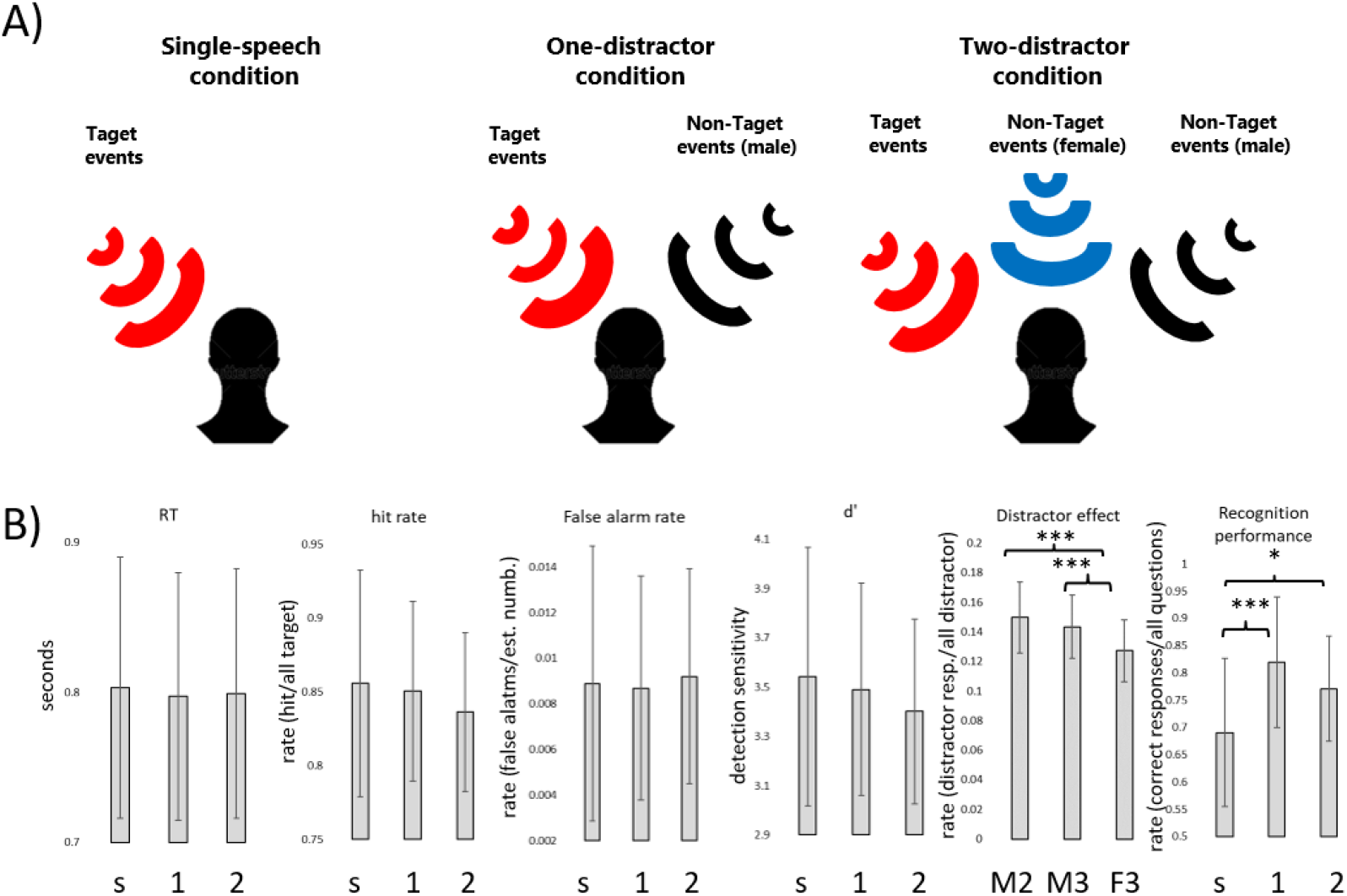
Top (A): Schematic illustration of the three experimental conditions. Participants were instructed to track the contents of the speech presented from the target location while also performing a numeral detection task on the same stream. In separate experimental conditions, participants were presented with 1) one speech stream (single-speech condition), 2) two speech streams (one-distractor condition), or 3) three speech streams (two-distractor condition). The target stream was always presented from the left of the listener (red loudspeaker). The black and blue wave symbols indicate the distractor male and distractor female streams, respectively. Bottom (B): Group average (N=23) performance (mean and standard deviation) in the numeral detection task indexed by RT, hit rate, false alarm rate, d’, and the distractor effect, as well as recognition performance in the content-tracking task, separately for the three experimental conditions (s: single-speech condition, 1: one-distractor condition, 2: two-distractor condition). Note that because no distractor was present in the single-stream condition, the distractor effect was only calculated for the male speaker in the one-distractor (M2), the male speaker in the two-distractor (M3), and the female speaker in the two-distractor (F3) condition. “*” = p<.05, “***” p<.001, and “+” = .05<p<.1

Each article contained 45-57 numerals (M=50.7, SD=2.7) of 2-4 syllable length. These served as targets in the target stream (targets) and distractors in the distractor stream(s). Only numerals indicating the quantity of some object within the context of the text were assigned as targets/distractors. For example, in Hungarian, the indefinite article (“egy”) is the same as the word for “one”. This word, when used as an article, did not constitute a target/distractor. There are also words, such as the Hungarian word for moonflower or daisy (“százszorszép” – literally translated as “hundred-times-beautiful”), which have a numeral as a component. These were not regarded as targets/distractors either. The temporal separation between successive target and distractor events was not controlled, because the articles serving as target and distractor streams were randomly paired, separately for each participant. In a representative example, the mean difference was calculated between target and distractor events: the mean difference was 2.348 s (SD: 1.722 s, min: 0.013 s, max: 7.535 s). Distractor articles (but targets not) also contained 19-26 syntactic violations (M=20.5, SD=1.4), which served for control purposes, as Szalárdy et al. (2018, 2019; 2020) found that when participants follow one of two concurrent speech streams, syntactic violations within the unattended stream do not elicit the syntax-violation related ERP components. Therefore, syntactic violations could be used to indicate whether the non-target stream(s) were attended or not.

Altogether 12 stimulus blocks were created from the 24 articles, and delivered in three stimulus conditions: 1) the Single-speech, 2) the One-distractor, and 3) the Two-distractor condition. Each condition received 4 stimulus blocks.

### Experimental procedure

The study was conducted in an acoustically attenuated, electrically shielded, dimly lit room at the Research Centre for Natural Sciences, Budapest, Hungary. Three Mackie MR5 mk3 Powered Studio Monitor loudspeakers were placed at an equal 200 cm distance from the participant, positioned symmetrically at − 30° (left) 0° (middle), and 30° (right) from the midline. Additionally, a 23” monitor was placed at 195 cm in front of the participant, showing an unchanging fixation cross (“+”) during the stimulus blocks. Participants were instructed to avoid eye blinks and other muscle movements and to watch the fixation cross while listening to the speech segments. EEG was recorded during the experimental blocks.

Participants performed two tasks on the target speech segments (Fig. 1): the “numeral detection” and the “content tracking” task. In the numeral detection task, participants were instructed to press a hand-held response key with their right thumb as soon as they detected the presence of a numeral word (target events, see above). For the content tracking task, they were informed that at the end of each stimulus block, they will have to answer 5 questions regarding the contents of the target speech segment. Each question corresponded to a piece of information that appeared within the target speech segment. The experimenter read the question and the 4 possible answers. The listener was then asked to verbally indicate his/her choice for the correct answer (multiple-choice test). The experimenter noted the participant’s choice and followed up with a request for the participant to assess his/her confidence for the choice from four alternatives: “I don’t remember I was just guessing”, “I am not sure, but the option I chose sounded familiar: I think I heard it during the last block”, “I am sure; I remember having heard it during the last block”, “I know the answer from some other source”. The confidence judgment was recorded by the experimenter. The two concurrent tasks served complementary purposes in directing the listener’s attention: Whereas the tracking task required listeners to integrate information over longer periods of time and to fully process the target speech segments, the detection task ensured that attention was continuously focused on the target speech segment.

The stimulus blocks were presented in pseudorandomized order: in the first half of the experimental session (blocks 1-6), each condition was presented two times in random order with the restriction that no condition was immediately repeated; in the second half of the session (blocks 7-12), conditions were presented in reversed order with respect to the first half. Participants were allowed to take a break during the experiment after each stimulus block, and there was a longer mandatory break after the 6th stimulus block. Altogether, the experiment lasted ca. 4 hours.

### Behavioral data recording and analysis

Detection task performance: Button presses for correct responses (hits) were initially collected from a 0–5000 ms interval from the onset of the target event. Responses were then rejected if they were longer than 95 % (>1493 ms) or shorter than 5 % (<435 ms) of all of the initially collected potential target responses (collapsed across all conditions and participants). From the accepted responses, log-normalized reaction times (RT) and hit rates (HR) were calculated for each participant and condition (pooling data from the four stimulus blocks of the same condition). False alarm rates (FA) were calculated by dividing the number of non-target responses (any response outside the periods calculated for targets) by the estimated number of non-target words in the target-stream (calculated from the mean word length for all speech material used in the experiment). Detection sensitivity values (d’; Green & Swets, 1966) were calculated from HR and FA. The distractor effect was assessed for distractor numerals: the number of distractors with a button press response (from the same time-interval as was found for the corresponding targets) was divided by the number of all distractors, separately for each condition and distractor source (one-distractor male, two-distractor male, two-distractor female).

Recognition performance in the content-tracking task was calculated as the percentage of correct responses, separately for each participant and condition (pooling data from the four stimulus blocks of the same condition). The sensitivity of the measurement was increased by eliminating items (questions), the response to which was above 95% or below 30% correct overall (collapsed across participants and conditions). Correct responses with a confidence judgment of “I know the answer from some other source” were also dropped from the analysis. Confidence ratings were calculated as responses “I don’t remember I was just guessing” received 1 point, “I am not sure, but the option I chose sounded familiar: I think I heard it during the last block” received 2 points, and “I am sure; I remember having heard it during the last block” received 3 points.

Statistical analysis consisted of repeated-measures analyses of variance (ANOVA) with the factor of CONDITION (single-speech vs. one-distractor vs. two-distractor), separately for RT, d’, hit rate, false alarm rate, and recognition performance. Statistical analysis of distractor effect (assessed for distractor numerals, see Methods) was performed by another repeated-measures ANOVA, with the factors DISTRACTOR (one-distractor male, two-distractor male, two-distractor female). The alpha level was set at 0.05. Greenhouse-Geisser correction of sphericity violations was employed where applicable and the ε correction factor is reported. All significant results are reported together with the η^2^ effect size. To evaluate the differences in the confidence rating between the three conditions (single-speech, one-distractor, two-distractor) Kruskal-Wallis H test was used.

All statistical analyses (behavioral and ERP) were conducted by the STATISTICA 13.1 and JASP 0.15.0.0. software.

### EEG data recording and analysis

EEG recording and analysis were identical to Szalárdy et al. (2018, 2019; 2020). Continuous EEG was recorded (1 kHz sampling rate and 100 Hz online low-pass filter) from a few seconds before the beginning to a few seconds after the end of the stimulus blocks using a BrainAmp DC 64-channel EEG system with actiCAP active electrodes (Brain Products GmbH). EEG signals were synchronized with the speech segments by matching an event trigger marked on the EEG record to the concurrent presentation of a beep sound in the audio stream (1 s before the speech segment commenced) with <1 ms accuracy. Electrodes were attached according to the extended International 10/20 system with an additional electrode placed on the tip of the nose. For identifying eye-movement artifacts, two electrodes were placed lateral to the outer canthi of the two eyes. Electrode impedances were kept below 15 kΩ. The FCz electrode served as the online reference.

Continuous EEG data were filtered with a 0.5-80.0 Hz Kaiser bandpass-filter and a 47.0-53.0 Hz Kaiser bandstop filter (the latter for removing electric noise; Kaiser β=5.65, filter length 18112 points) using the EEGlab 14.1.2.b toolbox (Delorme et al., 2007). EEG data processing was performed by Matlab R2018b (Mathworks Inc.). Electrodes showing long continuous or a large number of transient artifacts were substituted using the spline interpolation algorithm implemented in EEGlab. The maximum number of interpolated channels was two per participant. The Infomax algorithm of Independent Component Analysis (ICA) implemented in EEGlab was employed for eye-movement artifact removal. Maximum 6 ICA components (approximately 10 % of all components) constituting blink artifacts and horizontal eye-movements were removed via visual inspection of the topographical distribution and frequency contents of the components. Data were then offline re-referenced to the electrode attached to the tip of the nose. Epochs were extracted from continuous EEG records for a window of −200-2400 ms with respect to the onset of numerals. Baseline correction was applied using the 200-ms pre-event interval. Artifact rejection with a threshold of +/−100 μV voltage change was applied to the whole epoch, separately for each electrode. Artifact-free epochs were then averaged separately for each participant and condition. For target events, only hits, for distractors, only correct rejections were analyzed.

Amplitudes were measured from F3, Fz, F4, C3, Cz, C4, P3, Pz, P4 for statistical analysis. Time windows for measuring the ERP amplitudes were selected between 150 and 280 ms for N2b and between 620 and 770 ms for P3b relative to stimulus onset, both for target and non-target numerals. The average number of artifact-free target numerals were 150.17 (SD: 24.82) for the single-speech, 156.13 (SD: 24.77) for the one-distractor, and 160.39 (SD: 23.30) for the two-distractor condition. For distractor numerals and syntactic violations the average number of artifact-free trials were 179.26 (numeral, SD:24.77) and 74.13 (syntactic violation, SD: 9.09) for the one-distractor condition, 171.17 (numeral, SD: 21.92) and 72.52 (syntactic violation, SD: 8.81) for the two-distractor male, and 184.26 (numeral, SD: 21.26) and 72.30 (syntactic violation, SD: 8.88) for the two-distractor female speaker.

Target ERP amplitudes were statistically analyzed using repeated-measures ANOVAs with the factors of CONDITION (single-speech [“s”], one-distractor [“1”], two-distractor [“2”]) × ANTERIOR-POSTERIOR (frontal, central, parietal) × LATERALITY (left, middle, right), separately for the N2b and P3b components. Distractor ERPs were analyzed similarly, using repeated-measures ANOVAs with the factors of DISTRACTOR (one–distractor/male [“M2”], two-distractor/male [“M3”], two-distractor/female [“F3”]) × ANTERIOR-POSTERIOR (frontal, central, parietal) × LATERALITY (left, middle, right), separately for the N2b and P3b components. Post-hoc tests were conducted for all main effects and interactions that included the CONDITION (for targets) or the DISTRACTOR (for distractors) factor by Tukey’s HSD. Greenhouse-Geisser correction of sphericity violations was employed where applicable and the ε correction factor is reported together with the η^2^ effect size. Only significant effects including the CONDITION/DISTRCATOR factor are reported in the main text (see the Supplement for tables of all ANOVA effects). ERP amplitudes elicited by the syntactic violations were compared to zero by one-tailed Student’s *t*-test, applying Bonferroni correction for multiple comparisons.

## Results

### Behavioral measures

The descriptive statistics of the behavioral performance are shown in Figure 1B. A significant effect of condition was found for recognition performance (F(2,44) = 12.246, n_p_^2^ = .358, ε = .798, p < .001). This was due to the significantly lower memory performance in the single-speech condition compared to both the one- (p < .001) and the two-distractor (p = .011) conditions, whereas the latter two did not significantly differ from each other (p = .161). No significant effects were found for the reaction times (p = .633), hit (p = .123) and false alarm rates (p = .676).

A significant effect of DISTRACTOR was found on the distraction effect (F(2,44) = 19.200, n_p_^2^ = .466, ε = .862, p < .001). The effect was caused by the significantly lower distractor effect of the numerals spoken by the female speaker (F3) in the two-distractor condition compared to the male speaker in both the one- and two-distractor conditions (M2, M3; p < .001, both). The latter two were not significantly different from each other (p = .225). The confidence judgment was significantly different between the three conditions (Chi square = 34.189, p < .001, df = 2). Post-hoc significance values were adjusted by the Bonferroni correction for multiple tests, showing that significantly larger confidence judgment occurred in the one-distractor condition compared to the single-speech and two-distractor conditions (p < .001, both) whereas these were not different from each other (p = 1.00).

### ERP measures

ERPs measured at the Pz electrode are shown on Figure 2. The scalp distributions of the target N2b and P3b show maximal amplitude for both components over parietal scalp locations (Figure 3), as was also seen in our previous studies (Szalárdy et al., 2018, 2019; Szalárdy, Tóth, Farkas, Hajdu, et al., 2020; Szalárdy, Tóth, Farkas, Orosz, et al., 2020a).

**Figure 2.**
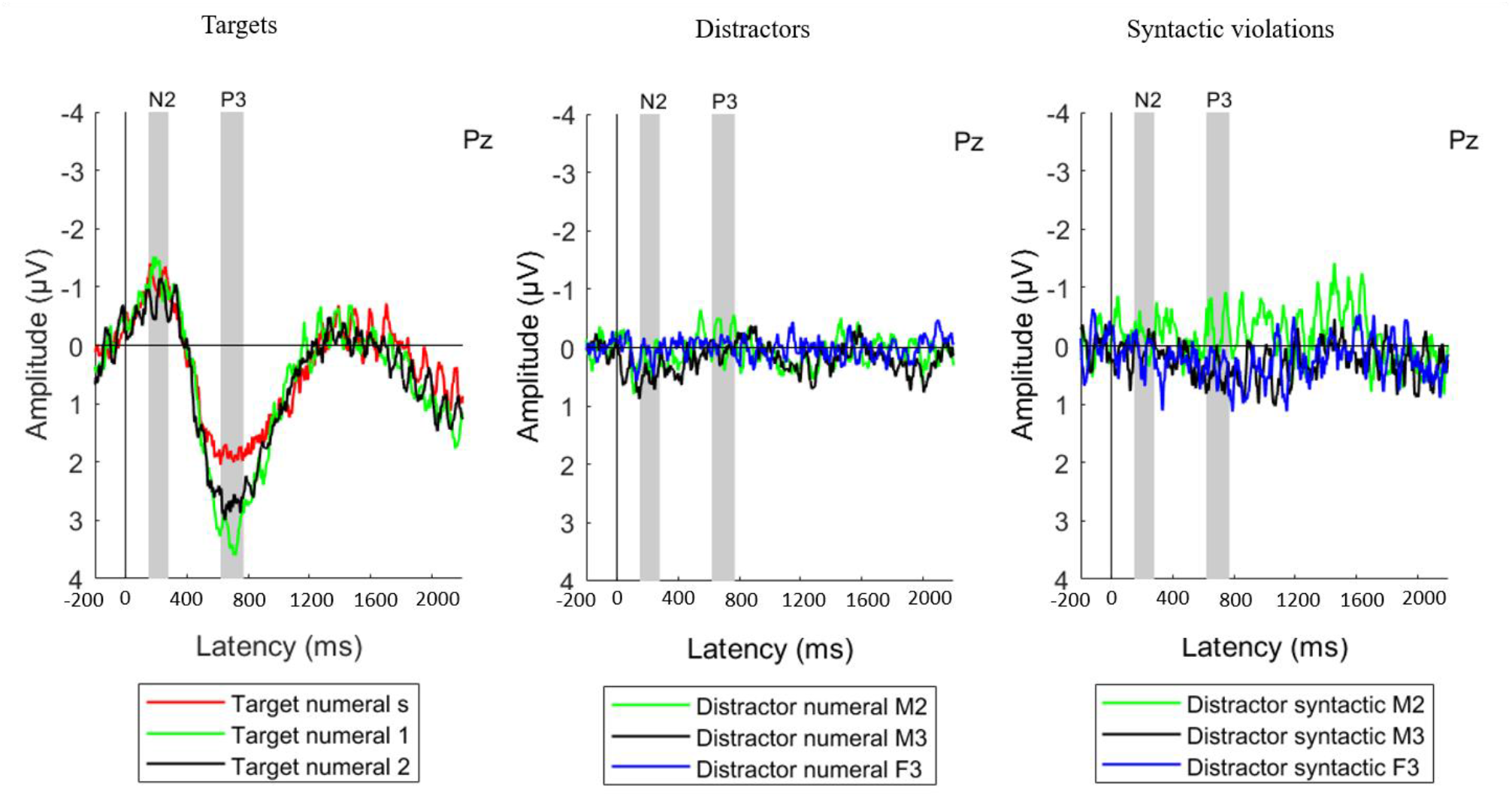
Group average (N=23) ERP responses measured from the parietal (Pz) electrode position, separately for the target numerals (left), distractor numerals (middle), and distractor syntactic violations (right). The measurement time windows for N2b and P3b are marked by grey vertical bands. Legend abbreviations for the target numerals: s – single-speech condition, 1 – one-distractor condition, 2 – two-distractor condition. Legend abbreviations for the distractor numerals and syntactic violations: M2 - male speaker in the one-distractor condition, M3 - male speaker in the two-distractor condition, F3-female speaker in the two-distractor condition.

**Figure 3.**
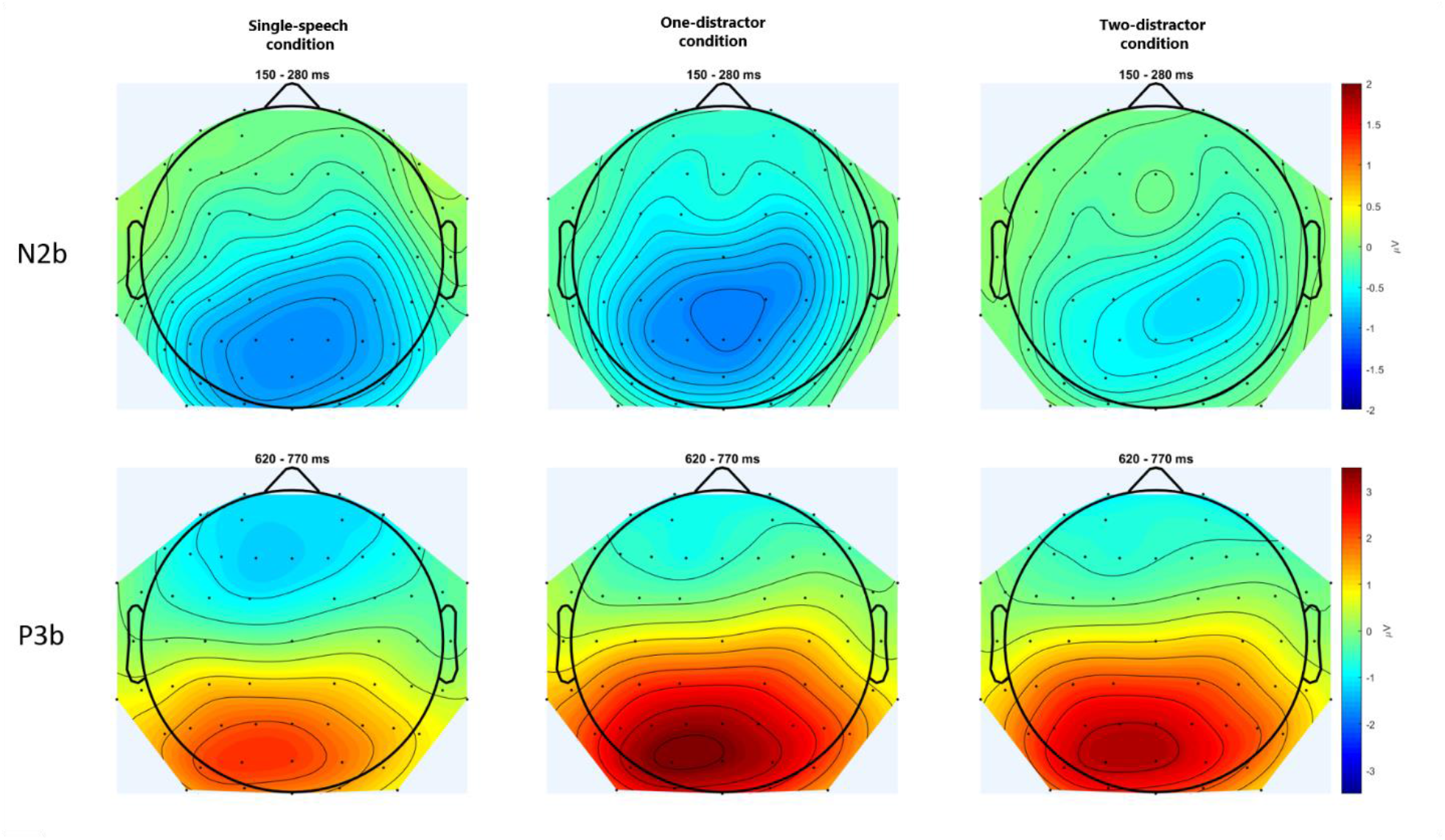
Scalp topography of the N2b (upper row) and P3b (bottom row) component for target numerals in the three experimental conditions: single-speech (left), one-distractor (middle), and two-distractor (right). The scalp distributions were calculated from the average voltages measured from the time windows shown in Figure 2. Maps were spline interpolated with a smoothing factor of 10^−7^.

#### ERPs to targets

For target events, significant interaction was found for the N2b amplitude between CONDITION and LATERALITY (F(2,44) = 2.862; ε = .799 p = .0398; ηp2 = .115). Post-hoc tests showed that the N2b amplitude significantly differed between all three conditions (s, 1, 2) on the left side (p = .0351, at least): the largest N2b amplitude was observed for “1” which decreased for “s” with the lowest amplitude for “2”. In contrast, the N2 amplitudes for “s” and “1” were not significantly different from each other at the midline (p = .88) or on the right side (p = 1.00) with those for “2” significantly differing from both at the midline (p < .001, both). In all of these cases, the amplitude for the target N2b was lower for “2” than for “s” and “1”. A significant main effect of CONDITION was found for the P3b component (F(2,44) = 18.580; ε = .987 p < .001; ηp2 = .458) with interactions between CONDITION and LATERALITY (F(4,88) = 3.973; ε = .827 p = .009; ηp2 = .153), and CONDITION, LATERALITY, and ANTERIOR-POSTERIOR (F(8,176) = 2.575; ε = .638 p = .029; ηp2 = .105). As P3b is maximal over parietal sites, for post-hoc analysis, a separate ANOVA was conducted on the parietal line, alone, with the factors of CONDITION and LATERALTY. In this post-hoc analysis, main effects of CONDITION (F(2,44) = 15.956; ε = .859 p < .001; ηp2 = .420) and LATERALITY (F(2,44) = 13.921; ε = .763 p < .001; ηp2 = .388) were found with no interaction between them (p = .292). The post-hoc test of the CONDITION main effect showed significantly lower amplitudes for “s” compared to “1” and “2” (p < .001, both), while the latter two were not significantly different from each other (p = .204).

Based on the similar pattern between the recognition performance data and P3b amplitude in the three conditions, Pearson correlation was calculated between them, using the P3b measured at the Pz electrode. No significant correlation was found between the P3b amplitude and recognition performance in the single stream (r = −.009; p = .966), one-distractor (r = −.112; p = .621), and two-distractor conditons (r = −.376; p = .077).

#### ERPs to distractors and syntactic violations

The positive deflection measured for the distractors in the N2b time window (Figure 2, middle; Figure 4 for scalp distributions), a significant DISTRACTOR × ANTERIOR-POSTERIOR interaction was found (F(4,88) = 3.429; ε = .547 p = .037; ηp2 = .135). Post-hoc tests revealed that this interaction was due to the central ERP amplitude in the N2b time window for “M3” being more positive than that for “M2” centrally (p = .033), and also than “M2” and “F3” parietally (p = .009, both). In contrast, no significant difference was found for the amplitudes from the N2b window between the “M2” and “F3” either over central (p = .491) or parietal electrode locations (p = 1.000). No other significant difference was found either in the N2b or the P3b time window.

**Figure 4.**
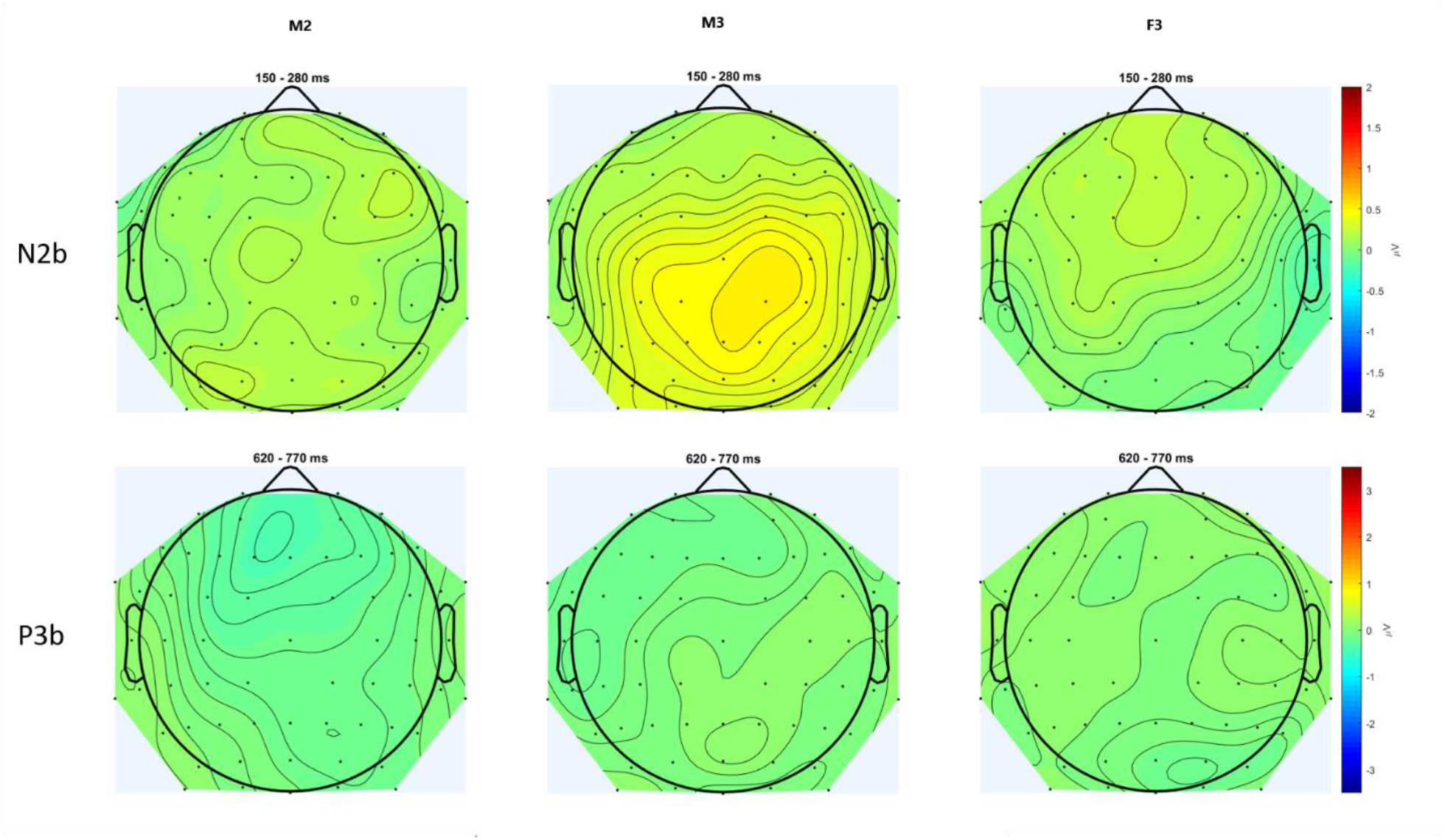
Scalp topography of the N2b (upper row) and P3b (bottom row) component for distractor numerals for the three distractor events: M2 - male speaker in the one-distractor condition, M3 - male speaker in the one-distractor condition, F3-female speaker in the two-distractor condition. The scalp distributions were calculated from the average voltages measured from the time windows shown in Figure 2. Maps were spline interpolated with a smoothing factor of 10^−7^.

Finally, syntactic violations did not elicit significant ERP components in the N2b or the P3b time window (Figure 2, right panel; p > .274, at least).

## Discussion

We investigated whether multiple distractor speech streams are segregated from each other and their effects on the lexical/semantical processing of the target speech stream. The current data corroborated previous findings (Bronkhorst, 2015) in that detection sensitivity and the target N2b amplitude decreased whereas the P3b amplitude increased from the single speech to the concurrent speech streams conditions. Importantly, the behavioral distractor effect differed between the conditions with one vs. two distractors (distraction by the female speaker was lower than that of the male speaker in either condition) and the target N2b elicited in the presence of two distractors was significantly smaller than that elicited in the presence of one distractor. Further, the positive deflection in the N2b time window significantly differed between the one- and two-distractor conditions for the same (M2) speaker. These results show that speech processing was different in the presence of one vs. two distractors, and thus, in terms of the alternatives described in the introduction, the current data suggest that the two background speech streams were segregated from each other.

Based on the behavioral results, the smaller distracting effect of the female voice alone may be explained in the context of both alternatives. It is compatible with the notion of segregating the two background streams with the additional assumption that distractors in the female voice were less likely to be confused with the target spoken in a male voice than those of another male voice. However, one could also assume that within the undifferentiated background, the female voice was a less effective masker for the target stream. Previous studies showed that target detection is reduced when a speech segment is masked by high-level noise (energetic masking), whereas allocation problems can be found when a speech stream is masked by another speech (informational masking; Darwin, 2008; Ihlefeld & Shinn-Cunningham, 2008). The current results showed no significant hit rate or false alarm difference between the one- and two-distractor conditions. Because the male distractor was common to both conditions, the lack of significant change in task performance is compatible with the less effective masker explanation.

The undifferentiated background hypothesis is, however, contrasted by both the target N2b and the distractor ERPs measured at the N2b window. The former is incompatible with the above suggested non-significant behavioral target-detection performance effect of the addition of the female distractor stream on top of the male masker stream. The N2b is a target-related response that has been associated with stimulus classification (Näätänen, 1990; Ritter et al., 1979) and its amplitude is modulated by selective attention (Michie, 1984). In contrast to our hypotheses, we found that the amplitude of the N2b did not linearly increase with task demand, but increased for the one-distractor conditions and decreased for the two-distractor condition. The decrease of the target N2b amplitude suggests that the identification of targets (the assumed role of the processes reflected by N2b - Näätänen, 1990; Ritter et al., 1979) differed between the one-distractor and two-distractor conditions. The two conditions were different in the background streams only, and the target properties were identical. The distractor stream thus served as a masker, which energetically and informationally masked the target stream. If masking (whether energetic or information) was the only way target identification was affected, then the N2b amplitude should have corresponded to the behavioral effects, mirroring the lack of difference found for P3b. Furthermore, in a previous study, the N2b was not sensitive to the effect of informational and energetic masking, but rather, the amplitude was modulated by attention (Szalárdy et al., 2019). The significant N2b difference observed may be explained by attentional selection: assuming that in a multi-stream situation, target detection must also include validating targets by taking into account the stream the candidate belongs to.

The strongest evidence supporting the segregation of the two background streams comes from the increased positivity in the N2b time window for the same distractor male speaker in the presence vs. the absence of the stream delivered by the female speaker. This suggests that distractors are processed differently alone than in the presence of another distractor stream. If we assume that the responses in the N2b latency range reflect target identification processes, then the differential response to the same distractor between the one- and the two-distractor condition reflects that target identification (rejection of the distractor) within the male distractor stream proceeded under a higher processing load due to the presence of the second distractor stream (i.e., the target stream was present in both conditions). The presence of another distractor results in higher information density and thus the allocation of reduced capacities to each stream, which in turn modulates both the target N2b (as discussed before) and the processes in N2b range of the distractors (see Conroy & Polich, 2007; Dowling et al., 2008; Isreal et al., 1980; Szalárdy et al., 2019). Crucially, identifying and rejecting target candidates in a distractor stream requires the stream to be segregated from both the target and the other distractor stream. Therefore, the results support the notion of segregating the background in the current situation. This conclusion is compatible with models suggesting full object-based description of the environment (for a general model of learning, see, Fiser, 2009; in the auditory modality, see e.g., Mill et al., 2013; Winkler et al., 2012).

However, the current data do not prove that the background is always parsed into its constituent streams. There are studies showing that a background consisting of potentially separable streams remained undistinguished (e.g., Brochard et al., 1999; Sussman et al., 2005). The crucial difference between these and the current study is the type of sounds presented in the different streams. While the studies found no segregation of streams in the background delivered simple sounds (mainly pure tones) differing from each other in one feature, here we presented natural speech, and specifically, the two unattended streams differed in the speaker’s gender, making them highly distinctive. In a recent study, attended and ignored speech streams were both represented in the auditory cortex (mostly in primary areas), suggesting the global representation of the full auditory scene with all auditory streams (Puvvada & Simon, 2017). Other studies also confirmed the recognition of some words from a background speech stream, even if the background consisted of multiple voices (Dekerle et al., 2014). Furthermore, signs of spectro-temporal and linguistic processing of task-irrelevant speech streams were found in the auditory cortex, left inferior cortex, and posterior parietal cortex (Brodbeck et al., 2020; Har-shai Yahav & Zion Golumbic, 2021). The prerequisite of background stream segregation might be highly distinctive features, which results in categorical differences, such as different gender of speakers; but this background stream segregation might be unique to speech perception. Cusack and colleagues (2004) have already speculated that distinctive acoustic features could allow streams to be segregated outside the focus of attention, and several studies have shown stream segregation when none of the streams was specifically attended (e.g., Sussman, 2007; Sussman et al., 2005). It is, therefore, possible that segregation of the background depends on both the perceptual difficulty of the separation (Cusack et al., 2004) and the available capacity (Sussman et al., 2005). This question requires future studies to explain the prerequisite of background stream segregation and whether it is specific to speech streams or other sounds as well.

Somewhat surprisingly, recognition performance was significantly lower in the single-speech condition compared to the conditions with distractors. This pattern of results was accompanied by a corresponding P3b amplitude effect, and the N2b was also lower for this condition suggesting poorer allocation of attention.

Furthermore, the confidence judgment was also lower in the single-speech condition compared to the one-distractor condition, but not in the two-distractor condition. Similar correspondence was found between recognition performance and the P3b amplitude in our previous experiment based on similar methods but presenting only a single distractor (Szalárdy et al., 2019). This is not a trivial finding, as P3b was elicited in a task (numeral detection) concurrent to the memory task (which was only tested after the stimulus blocks). Studies testing working memory also found that better performance was associated with higher P3b amplitude, especially with higher motivational salience (e.g., reward, punishment; Baskin-Sommers et al., 2014) and for young healthy adults (Saliasi et al., 2013). Thus, it is possible that performing the tasks under more difficult circumstances (in the presence of distractor stream(s)) resulted in better engagement with the task, which boosted performance in content tracking. Alternatively, performing the target detection task at a high level in the presence of distractor streams forced participants to utilize higher-level speech cues (syntactic and semantic) in order to determine whether a given numeral (candidate target) indeed needed a response. Investing more effort in fully processing the target speech stream could have resulted in better memory for the contents of the speech stream, and thus higher recognition performance. Although the current results do not allow us to separate the alternative explanations, they corroborate the previously observed correspondence between recognition memory performance and the P3b amplitude.

In summary, the present data support the parsing of the background into separate speech streams, based on the pattern of behavioral and ERP effects obtained for both target and distractor events. Together they suggest that lexical-semantic analysis operated to some degree also on the distractor speech streams, requiring them to be segregated from each other as well as from the target stream. The data also showed parallel effects of the manipulations on the P3b amplitude to a numeral detection task and recognition memory performance (tested after the stimulus blocks), which may be commonly governed by the level of task engagement or by the depth of speech processing necessitated by auditory stream segregation.

## Acknowledgment

This work was funded by the National Research, Development and Innovation Office (project K132642). The authors are grateful to Zsuzsanna D'Albini and Zsuzsanna Kovács for collecting the EEG data, Ágnes Palotás and László Hunyadi for text editing, to László Liszkai for audio recording and editing, to Zsuzsanna Kocsis for the ICA data preprocessing, to Gábor Urbán, Botond Hajdu and Bálint File for providing help in the analysis scripts, and to Ferenc Elek and Péter Scherer for voicing the articles.

## References

Astheimer, L. B., & Sanders, L. D. (2009). Listeners modulate temporally selective attention during natural speech processing. Biological Psychology, 80(1), 23–34. https://doi.org/10.1016/j.biopsycho.2008.01.015

Baskin-Sommers, A. R., Krusemark, E. A., Curtin, J. J., Lee, C., Vujnovich, A., & Newman, J. P. (2014). The impact of cognitive control, incentives, and working memory load on the P3 responses of externalizing prisoners. Biological Psychology, 96(1), 86–93. https://doi.org/10.1016/j.biopsycho.2013.12.005

Best, V., Ozmeral, E. J., Kopčo, N., & Shinn-Cunningham, B. G. (2008). Object continuity enhances selective auditory attention. Proceedings of the National Academy of Sciences of the United States of America, 105(35), 13174–13178. https://doi.org/10.1073/pnas.0803718105

Bregman, A. S. (1990). Auditory scene analysis: The perceptual organization of sound (A. S. Bregman (ed.)). The MIT Press. https://doi.org/10.7551/mitpress/1486.001.0001

Brochard, R., Drake, C., Botte, M. C., & McAdams, S. (1999). Perceptual organization of complex auditory sequences: Effect of number of simultaneous subsequences and frequency separation. Journal of Experimental Psychology: Human Perception and Performance, 25(6), 1742–1759. https://doi.org/10.1037/0096-1523.25.6.1742

Brodbeck, C., Jiao, A., Hong, L. E., & Simon, J. Z. (2020). Neural speech restoration at the cocktail party: Auditory cortex recovers masked speech of both attended and ignored speakers. PLoS Biology, 18(10). https://doi.org/10.1371/JOURNAL.PBIO.3000883

Bronkhorst, A. W. (2015). The cocktail-party problem revisited: early processing and selection of multi-talker speech. Attention, Perception, and Psychophysics, 77(5), 1465–1487. https://doi.org/10.3758/s13414-015-0882-9

Cherry, E. C. (1953). Some experiments on the recognition of speech, with one and with two ears. Journal of the Acoustical Society of America, 25(5), 975–979. https://doi.org/10.1121/1.1907229

Conroy, M. A., & Polich, J. (2007). Normative variation of P3a and P3b from a large sample: Gender, topography, and response time. Journal of Psychophysiology, 21(1), 22–32. https://doi.org/10.1027/0269-8803.21.1.22

Cusack, R., Deeks, J., Aikman, G., & Carlyon, R. P. (2004). Effects of location, frequency region, and time course of selective attention on auditory scene analysis. Journal of Experimental Psychology: Human Perception and Performance, 30(4), 643–656. https://doi.org/10.1037/0096-1523.30.4.643

Darwin, C. J. (2008). Listening to speech in the presence of other sounds. Philosophical Transactions of the Royal Society B: Biological Sciences, 363(1493), 1011–1021. https://doi.org/10.1098/rstb.2007.2156

Dekerle, M., Boulenger, V., Hoen, M., & Meunier, F. (2014). Multi-talker background and semantic priming effect. Frontiers in Human Neuroscience, 8(October), 1–13. https://doi.org/10.3389/FNHUM.2014.00878/ABSTRACT

Delorme, A., Sejnowski, T., & Makeig, S. (2007). Enhanced detection of artifacts in EEG data using higher-order statistics and independent component analysis. NeuroImage, 34(4), 1443–1449. https://doi.org/10.1016/j.neuroimage.2006.11.004

Donchin, E., & Coles, M. G. H. (1988). Is the P300 component a manifestation of context updating? Behavioral and Brain Sciences, 11(3), 357–374. https://doi.org/10.1017/S0140525X00058027

Dowling, W. J., Bartlett, J. C., Halpern, A. R., & Andrews, M. W. (2008). Melody recognition at fast and slow tempos: Effects of age, experience, and familiarity. Perception and Psychophysics, 70(3), 496–502. https://doi.org/10.3758/PP.70.3.496

Fiser, J. (2009). Perceptual learning and representational learning in humans and animals. Learning & Behavior, 37(2), 141–153. https://doi.org/10.3758/LB.37.2.141

Fitzgerald, K., & Todd, J. (2020). Making Sense of Mismatch Negativity. Frontiers in Psychiatry, 11, 468. https://doi.org/10.3389/fpsyt.2020.00468

Green, D. M., & Swets, J. A. (1966). Signal detection theory and psychophysics. Wiley.

Har-shai Yahav, P., & Zion Golumbic, E. (2021). Linguistic processing of task-irrelevant speech at a cocktail party. ELife, 10. https://doi.org/10.7554/ELIFE.65096

Ihlefeld, A., & Shinn-Cunningham, B. (2008). Spatial release from energetic and informational masking in a divided speech identification task. The Journal of the Acoustical Society of America, 123(6), 4380–4392. https://doi.org/10.1121/1.2904825

Isreal, J. B., Chesney, G. L., Wickens, C. D., & Donchin, E. (1980). P300 and tracking difficulty: evidence for multiple resources in dual‐task performance. Psychophysiology, 17(3), 259–273. https://doi.org/10.1111/j.1469-8986.1980.tb00146.x

Kidd, G., Mason, C. R., & Gallun, F. J. (2005). Combining energetic and informational masking for speech identification. The Journal of the Acoustical Society of America, 118(2), 982–992. https://doi.org/10.1121/1.1953167

Lambrecht, J., Spring, D. K., & Münte, T. F. (2011). The focus of attention at the virtual cocktail party-Electrophysiological evidence. Neuroscience Letters, 489(1), 53–56. https://doi.org/10.1016/j.neulet.2010.11.066

Micheyl, C., Shamma, S. A., & Oxenham, A. J. (2007). Hearing Out Repeating Elements in Randomly Varying Multitone Sequences: A Case of Streaming? In Hearing – From Sensory Processing to Perception (pp. 267–274). Springer Berlin Heidelberg. https://doi.org/10.1007/978-3-540-73009-5_29

Michie, P. T. (1984). Selective sttention effects on somatosensory event‐related potentials. Annals of the New York Academy of Sciences, 425(1), 250–255. https://doi.org/10.1111/j.1749-6632.1984.tb23542.x

Mill, R. W., Bohm, T. M., Bendixen, A., Winkler, I., & Denham, S. L. (2013). Modelling the Emergence and Dynamics of Perceptual Organisation in Auditory Streaming. PLoS Computational Biology, 9(3), 1002925. https://doi.org/10.1371/journal.pcbi.1002925

Näätänen, R. (1990). The role of attention in auditory information processing as revealed by event-related potentials and other brain measures of cognitive function. Behavioral and Brain Sciences, 13(2), 201–233. https://doi.org/10.1017/S0140525X00078407

Näätänen, R., Simpson, M., & Loveless, N. E. (1982). Stimulus deviance and evoked potentials. Biological Psychology, 14(1–2), 53–98. https://doi.org/10.1016/0301-0511(82)90017-5

Nasman, V. T., & Rosenfeld, J. P. (1990). Parietal P3 response as an indicator of stimulus categorization: increased P3 amplitude to categorically deviant target and nontarget stimuli. Psychophysiology, 27(3), 338–350. https://doi.org/10.1111/j.1469-8986.1990.tb00393.x

Polich, J. (2003). Theoretical overview of P3a and P3b. In J. Polich (Ed.), Detection of change: event-related potential and fMRI findings (pp. 83–98). Kluwer Academic Press. https://doi.org/10.1007/978-1-4615-0294-4_5

Polich, J. (2007). Updating P300: An integrative theory of P3a and P3b. Clinical Neurophysiology, 118(10), 2128–2148. https://doi.org/10.1016/j.clinph.2007.04.019

Polich, J., & Herbst, K. L. (2000). P300 as a clinical assay: Rationale, evaluation, and findings. International Journal of Psychophysiology, 38(1), 3–19. https://doi.org/10.1016/S0167-8760(00)00127-6

Puvvada, K. C., & Simon, J. Z. (2017). Cortical Representations of Speech in a Multitalker Auditory Scene. Journal of Neuroscience, 37(38), 9189–9196. https://doi.org/10.1523/JNEUROSCI.0938-17.2017

Rahne, T., Böckmann, M., von Specht, H., & Sussman, E. S. (2007). Visual cues can modulate integration and segregation of objects in auditory scene analysis. Brain Research, 1144(1), 127–135. https://doi.org/10.1016/j.brainres.2007.01.074

Ritter, W., Simson, R., & Vaughan, H. G. (1983). Event‐related potential correlates of two stages of information processing in physical and semantic discrimination tasks. Psychophysiology, 20(2), 168–179. https://doi.org/10.1111/j.1469-8986.1983.tb03283.x

Ritter, W., Simson, R., Vaughen, H. G., & Friedman, D. (1979). A brain event related to the making of a sensory discrimination. Science, 203, 1358–1361. https://doi.org/10.1126/science.424760

Saliasi, E., Geerligs, L., Lorist, M. M., & Maurits, N. M. (2013). The Relationship between P3 Amplitude and Working Memory Performance Differs in Young and Older Adults. PLoS ONE, 8(5), e63701. https://doi.org/10.1371/journal.pone.0063701

Siegenthaler, B. M., & Barr, C. A. (1967). Auditory Figure-Background Perception in Normal Children. Child Development, 38(4), 1163. https://doi.org/10.2307/1127113

Snyder, J. S., & Alain, C. (2007). Toward a neurophysiological theory of auditory stream segregation. Psychological Bulletin, 133(5), 780–799. https://doi.org/10.1037/0033-2909.133.5.780

Southwell, R., Baumann, A., Gal, C., Barascud, N., Friston, K., & Chait, M. (2017). Is predictability salient? A study of attentional capture by auditory patterns. Philosophical Transactions of the Royal Society B: Biological Sciences, 372(1714). https://doi.org/10.1098/rstb.2016.0105

Sussman, E. S. (2007). A new view on the MMN and attention debate: The role of context in processing auditory events. Journal of Psychophysiology, 21(3–4), 164–175. https://doi.org/10.1027/0269-8803.21.34.164

Sussman, E. S., Bregman, A. S., Wang, W. J., & Khan, F. J. (2005). Attentional modulation of electrophysiological activity in auditory cortex for unattended sounds within multistream auditory environments. Cognitive, Affective and Behavioral Neuroscience, 5(1), 93–110. https://doi.org/10.3758/CABN.5.1.93

Sussman, E. S., Horváth, J., Winkler, I., & Orr, M. (2007). The role of attention in the formation of auditory streams. Perception and Psychophysics, 69(1), 136–152. https://doi.org/10.3758/BF03194460

Sussman, E. S., Ritter, W., & Vaughan, H. G. (1998). Attention affects the organization of auditory input associated with the mismatch negativity system. Brain Research, 789(1), 130–138. https://doi.org/10.1016/S0006-8993(97)01443-1

Szalárdy, O., Bőhm, T. M., Bendixen, A., & Winkler, I. (2013). Event-related potential correlates of sound organization: Early sensory and late cognitive effects. Biological Psychology, 93(1). https://doi.org/10.1016/j.biopsycho.2013.01.015

Szalárdy, O., Tóth, B., Farkas, D., György, E., & Winkler, I. (2019). Neuronal correlates of informational and energetic masking in the human brain in a multi-talker situation. Frontiers in Psychology, 10. https://doi.org/10.3389/fpsyg.2019.00786

Szalárdy, O., Tóth, B., Farkas, D., Hajdu, B., Orosz, G., & Winkler, I. (2020). Who said what? The effects of speech tempo on target detection and information extraction in a multi-talker situation: An ERP and functional connectivity study. Psychophysiology. https://doi.org/10.1111/psyp.13747

Szalárdy, O., Tóth, B., Farkas, D., Kovács, A., Urbán, G., Orosz, G., Szabó, B. T., Hunyadi, L., Hajdu, B., & Winkler, I. (2018). The effects of attention and task-relevance on the processing of syntactic violations during listening to two concurrent speech streams. Cognitive, Affective and Behavioral Neuroscience, 18(5). https://doi.org/10.3758/s13415-018-0614-4

Szalárdy, O., Tóth, B., Farkas, D., Orosz, G., Honbolygó, F., & Winkler, I. (2020a). Linguistic predictability influences auditory stimulus classification within two concurrent speech streams. Psychophysiology, 57(5). https://doi.org/10.1111/psyp.13547

Szalárdy, O., Tóth, B., Farkas, D., Orosz, G., Honbolygó, F., & Winkler, I. (2020b). Linguistic predictability influences auditory stimulus classification within two concurrent speech streams. Psychophysiology, 57(5). https://doi.org/10.1111/psyp.13547

Szalárdy, O., Winkler, I., Schröger, E., Widmann, A., & Bendixen, A. (2013). Foreground-background discrimination indicated by event-related brain potentials in a new auditory multistability paradigm. Psychophysiology, 50(12). https://doi.org/10.1111/psyp.12139

Teki, S., Chait, M., Kumar, S., Von Kriegstein, K., & Griffiths, T. D. (2011). Brain bases for auditory stimulus-driven figure-ground segregation. Journal of Neuroscience, 31(1), 164–171. https://doi.org/10.1523/JNEUROSCI.3788-10.2011

Tóth, B., Kocsis, Z., Háden, G. P., Szerafin, Á., Shinn-Cunningham, B. G., & Winkler, I. (2016). EEG signatures accompanying auditory figure-ground segregation. NeuroImage, 141, 108–119. https://doi.org/10.1016/j.neuroimage.2016.07.028

Wightman, F. L., & Kistler, D. J. (2005). Informational masking of speech in children: Effects of ipsilateral and contralateral distracters. The Journal of the Acoustical Society of America, 118(5), 3164–3176. https://doi.org/10.1121/1.2082567

Winkler, I., Denham, S., Mill, R., Bohm, T. M., & Bendixen, A. (2012). Multistability in auditory stream segregation: A predictive coding view. Philosophical Transactions of the Royal Society B: Biological Sciences, 367(1591), 1001–1012. https://doi.org/10.1098/rstb.2011.0359

Winkler, I., Van Zuijen, T. L., Sussman, E., Horváth, J., & Näätänen, R. (2006). Object representation in the human auditory system. European Journal of Neuroscience, 24(2), 625–634. https://doi.org/10.1111/j.1460-9568.2006.04925.x

Wood, N. L., & Cowan, N. (1995). The cocktail party phenomenon revisited: Attention and memory in the classic selective listening procedure of Cherry (1953). Journal of Experimental Psychology: General, 124(3), 243–262. https://doi.org/10.1037/0096-3445.124.3.243

Yuriko Santos Kawata, N., Hashimoto, T., & Kawashima, R. (2020). Neural mechanisms underlying concurrent listening of simultaneous speech. Brain Research, 1738, 146821. https://doi.org/10.1016/j.brainres.2020.146821

